# PIM-related kinases selectively regulate sensory functions in *C. elegans*

**DOI:** 10.1101/324046

**Authors:** Karunambigai S. Kalichamy, Kaisa Ikkala, Jonna Pörsti, Niina M. Santio, Sweta Jha, Carina I. Holmberg, Päivi J. Koskinen

## Abstract

The mammalian PIM family of serine/threonine kinases regulate several cellular functions, such as cell survival and motility. Since we have observed PIM expression in the olfactory epithelium and other sensory organs of mice, this has raised the question of whether PIM kinases regulate also sensory cell functions. As our model organism to investigate this question, we used the *Caenorhabditis elegans* nematodes, which express two PIM-related kinases, PRK-1 and PRK-2. We demonstrated them to be true PIM orthologs with similar substrate specificity as well as sensitivity to PIM-inhibitory compounds. Furthermore, we obtained evidence to indicate that PRKs are selectively involved in regulation of olfactory sensations via AWB or AWC^ON^ neurons to volatile attractants or repellants, but do not affect gustatory sensations.

## INTRODUCTION

Many organisms use chemosensation to detect beneficial and harmful substances in their living environment and to communicate with other animals. The nematode *C. elegans* is an ideal model to conduct genetic and behavioral studies on chemosensation, as it has a well-developed chemosensory system that enables it to discriminate between multiple kinds of volatile or water-soluble cues to find food and mating partners, and to avoid pathogens and other types of danger. *C. elegans* has a simple nervous system, which in the adult hermaphrodite is composed of 302 neurons with known synaptic connections. Among them, there are 22 amphid sensory neurons that fall into 11 morphological classes with one member of each class in the left and right side. A large number of volatile compounds can be sensed by *C. elegans* via three pairs of olfactory neurons, namely AWA, AWB and AWC neurons, whereas water-soluble compounds are sensed by the gustatory ASE and ASH neurons, which are responsive to both chemical and mechanical cues (Bargmann, 2006). The AWA, AWC and ASE neurons detect attractants, but AWB and ASH repellents. AWA neurons can detect odorants like 2,4,5-trimethylthiazole, diacetyl and pyrazine, while AWC neurons sense butanone, isoamyl alcohol (IAA), benzaldehyde, 2,3-pentanedione, and 2,4,5-trimethylthiazole (Bargmann et al., 1993). ASE neurons can detect soluble attractants like sodium or potassium chloride (Pierce-Shimomura et al., 2001). AWB neurons in turn react to volatile repellents like 2-nonanone or 1-octanol (Troemel et al., 1997), and ASH to soluble repellents like copper or SDS (Bargmann, 2006). The bilaterally symmetrical pairs of chemosensory neurons are structurally similar on each side of the animals. However, AWC and ASE amphid neurons are functionally distinct between the right and left sides and detect different compounds (Pierce-Shimomura et al., 2001; Wes and Bargmann, 2001; Yu et al., 1997).

In *C. elegans* neurons, the olfactory signals are received by G protein-coupled receptors that use cyclic GMP or polyunsaturated fatty acids as second messengers to open cation channels and to depolarize the neurons (Bargmann, 2006). These pathways may be modulated and fine-tuned by kinases and phosphatases. In the present study, we investigated the putative regulatory role of the serine/threonine-specific PIM-related kinases, PRKs. In mammalian cells, the oncogenic PIM family proteins (Brault et al., 2010; Nawijn et al., 2011; Santio and Koskinen, 2017) have been shown to phosphorylate multiple proteins such as the transcription factors NFATc1 (Rainio et al., 2002), RUNX1 and RUNX3 (Aho et al., 2006), NOTCH1 (Santio et al., 2016a), FOXP3 (Santio et al., 2016b), as well as the pro-apoptotic BAD protein (Aho et al., 2004; Yan et al., 2003) and the kinase GSK3B (Santio et al., 2016b) to enhance cell survival and motility, and thereby to promote metastatic growth of cancer cells (Santio et al., 2015). During mouse embryogenesis, the *pim* genes are mainly expressed in cells of the immune system and the central nervous system, including several sensory organs such as neural retina and olfactory epithelium (Eichmann et al., 2000). These observations have raised the questions of whether PIM kinases are required for the development of these organs and/or whether they regulate sensory cell functions.

In *C. elegans*, there are two homologs for the three mammalian PIM kinases, PRK-1 and PRK-2 (wormbase.org). PRK-1 isoforms are translated from four transcripts and PRK-2 from two. PRK-1 is expressed in both larvae and adults in the nervous system, intestine as well as head and tail neurons (wormbase.org). The expression pattern for PRK-2 is not known, but it has been reported to regulate neurite branching (Zheng et al., 2011), so most likely it is also expressed in neurons. Otherwise very little is known about the functions of PRKs. In this study, we used *in vitro* kinase assays to confirm that PRK-2 is a functional ortholog of mammalian PIM kinases. By using several PIM-selective inhibitors to investigate the roles of PRKs in chemosensation in *C. elegans*, we obtained evidence that they do not affect gustatory sensations, but selectively target AWC^ON^ and AWB neurons to regulate olfactory sensations.

## MATERIALS AND METHODS

### *C. elegans* culture and strains used

The *C. elegans* trains were grown and maintained on NGM agar plates seeded with the *E.coli* strain OP50, using standard culturing methods (Brenner, 1974). The wild-type Bristol N2 strain and the BC14881 (dpy-5(e907) I; *sEx14881*[rCesC06E8.3A∷GFP +pCeh361]) strain with GFP expression driven by the *prk-1* promoter (McKay et al., 2003) were obtained from the Caenorhabditis Genetics Center, MN, USA.

### Kinase assays

GST (glutathione S-transferase) fusion constructs of wild-type mouse PIM-1 and NFATc1 (Rainio et al., 2002) as well as *C. elegans* PRK-2 (a kind gift from Michael Nonet, Washington University, MO, USA) were used to produce proteins in bacteria. The proteins were purified with glutathione sepharose beads (GE Healthcare) and cleaved with PreScission protease (GE Healthcare) according to manufacturers instructions. To inhibit kinase activity, aliquots of PIM-1 or PRK-2 protein were preincubated for 10 min with the PIM-selective inhibitor DHPCC-9 (Akué-Gédu et al., 2009) that was dissolved in dimethyl sulfoxide (DMSO) at 10 μM concentration. 0.1% DMSO alone was used as a negative control. Radioactive kinase assays were carried out in the presence of gamma-^32^P-ATP (GE Healthcare) as previously described (Kiriazis et al., 2013). Phosphorylated proteins were resolved in 10% SDS-PAGE and stained with PageBlue^™^ solution (Thermo Fischer Scientific) to visualize protein loading. Radioactivity of the samples was analysed by autoradiography and the signal intensities were quantitated by Fiji (National Institutes of Health, Bethesda, MD, USA). Signals from phosphorylated samples were compared to the amounts of protein to calculate relative phosphorylation levels in the absence or presence of DHPCC-9.

### Kinase inhibitor treatments

To investigate the role of *Ce* PRKs in chemosensation, we used three structurally unrelated PIM-selective inhibitors, the pyrrolocarbazole carbaldehyde DHPCC-9 (Akué-Gédu et al., 2009), the imidazopyridazine SGI-1776 or the thiazolidinedione AZD-1208 (Medchem Express) that were all dissolved in DMSO. 0.1% DMSO alone was used as a negative control. Well-fed day one *C. elegans* adults synchronized at 20°C were exposed in S-basal buffer to various concentrations of the PIM inhibitors. After 120 min of exposure, animals were washed twice with the S-basal buffer and once with distilled water, and subjected to behavioral assays.

### Behavioral assays

#### Olfactory assays

The protocol for olfactory assays was adopted from Bargmann and coworkers (1993). The volatile odorants were diluted in ethanol, which was also used as the neutral control. The standard odorant dilutions used for chemotaxis assays were: butanone, 1:1000; benzaldehyde, 1:200; isoamyl alcohol, 1:100; 2,3-pentanedione, 1:10000; 2,4,5-trimethyl thiazole, 1:1000; diacetyl, 1:1000; pyrazine, 10 mg/ml and 1-octanol, 1:100. All odorants were purchased from Sigma-Aldrich, except for isoamyl alcohol from Fluka. 1 μl of 1M sodium azide (Sigma-Aldrich) was dropped in advance to both odorant and control spots to paralyze animals reaching them. Assay plates with animals were incubated for 120 min at 20°C, after which plates were moved to 4°C to stop animal movements. Chemotaxis indices (CI = O-C/O+C) were calculated as the number of animals that moved towards the attractive odorant (O) minus the number of animals towards the control (C), divided by the total number of animals. Avoidance indices (AI) were calculated similarly, but taken into account the number of animals that moved away from the repellant.

#### Gustatory assays

The protocol for gustatory assays with water-soluble attractants was adopted from Jansen and coworkers (2002) with slight modifications. Four quadrant lines were drawn on agar plates. 25 μl aliquots of 2.5 M NaCl or KCl (AnalaR Normapur) dissolved in distilled water were dropped on the opposite quadrants (A, C) and the solvent as control on the other pair of quadrants (B, D), after which the plates were allowed to dry at room temperature for 60 min. Control- and drug-exposed animals were washed and dropped in the middle of the assay plate and incubated for 60 min at 20°C. Then 0.5 μl of 1M sodium azide was added to the salt or solvent spots to paralyze the animals. Animals at each quadrant were counted and calculated for chemotaxis index (CI = (A+C) − (B+D) / A+B+C+D) as number of animals on the salt (A+C) minus number of animals on the solvent (B+D) divided by the total number of animals.

Aversion assays were performed according to Wicks and coworkers (2000) against water-soluble copper ions. 25 μl aliquots of 50 and 100 mM CuSO_4_ (Merck) solution were pipetted across the midlines of the assay plates. Approximately 200-250 animals were placed on one side and odorants or control spots on the other side. After 120 min of incubation at 20°C, chemotactic indices (CI = A/A+B) were calculated as the number of animals that had crossed the aversion compound line (A) divided by the total number of animals.

### Microscopy

Young day 4 adult animals of the *prk-1*::GFP strain were mounted on 3% agarose pads on glass slides and paralysed with 0.5 mM levamisole. For confocal imaging, a motorized Zeiss Axio Observer Z1 inverted microscope with Airyscan detector and Zeiss ZEN2 software was used. Images were taken with a 63x 1.4 NA plan-apochromat objective, using Argon laser with 488 nm wavelength.

### Statistical analysis

Data are presented as mean ± standard error of mean (SEM). All statistical analyses were performed using two-tailed paired student’s t-Test with equal variance. In each analysis, P<0.01 was used as a limit for significant difference.

## RESULTS

### Mammalian PIM kinases and *C. elegans* PIM-related kinases are true orthologs

An amino acid sequence alignment between the two *C. elegans* PIM-related kinases (PRKs) and the three human or mouse PIM kinases revealed that they all are well conserved with approximately 40% homology between the vertebrate and invertebrate proteins (Fig. 1A). Expectedly, the highest identities were observed within their kinase domains, especially at the ATP-binding site and the catalytically active site. By contrast, the C-terminal tails of *C. elegans* PRKs are much longer than those of their mammalian orthologs, and are not conserved even between PRK-1 and PRK-2.

**Figure 1.**
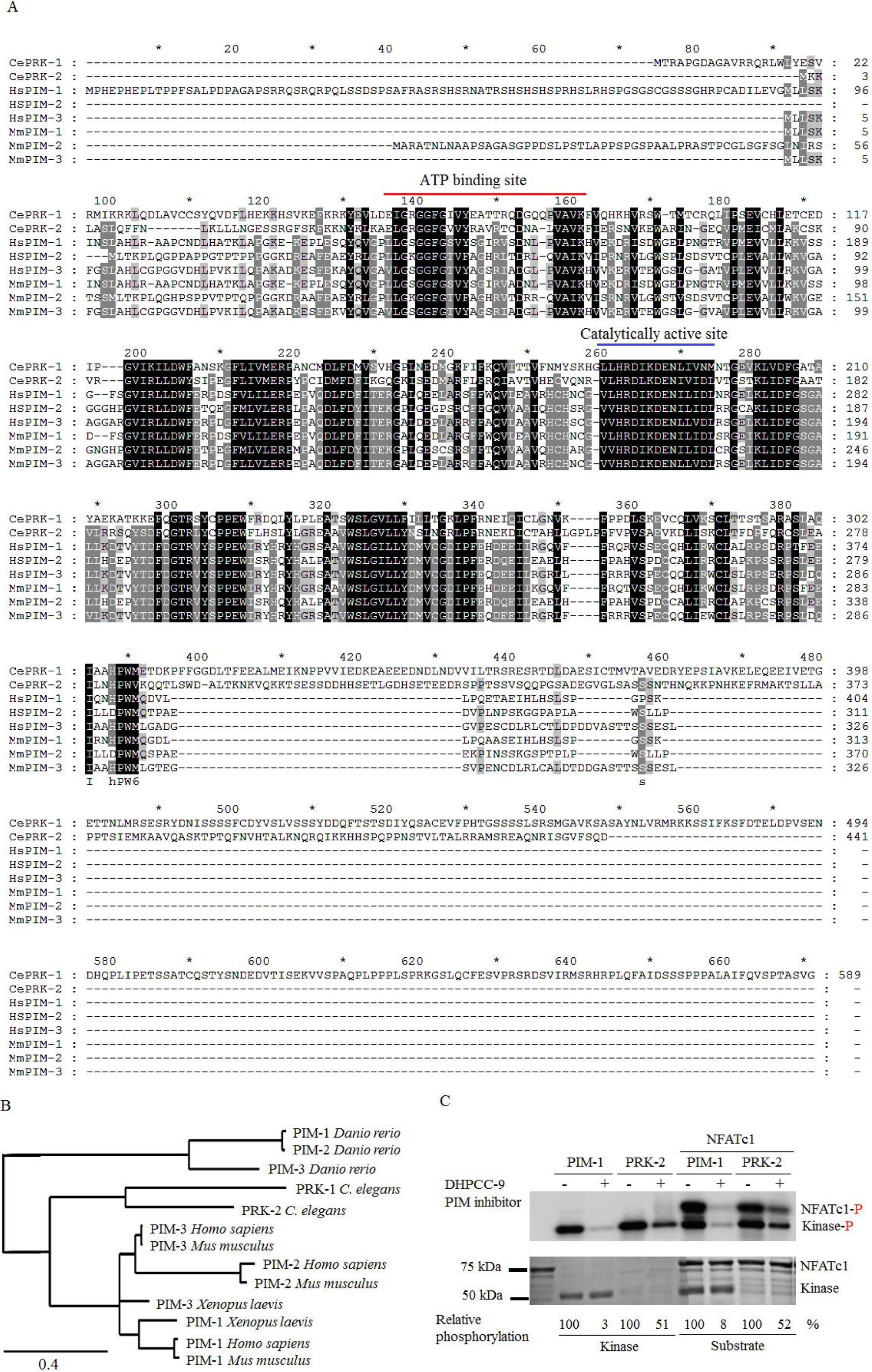
Mammalian PIM kinases and *C. elegans* PIM-related kinases are true orthologs. A. Amino acid sequences of *C. elegans* PRK-1 (NM_001276777.1) and PRK-2 (CAA84323.2) were aligned with *Homo sapiens* PIM-1 (NP_001230115.1), PIM-2 (NP_006866.2) and PIM-3 (NP_001001852.2), and *Mus musculus* PIM-1 (NP_032868.2), PIM-2 (NP_613072.1) and PIM-3 (NP_663453.1), using ClustalW. Identical or similar amino acids are marked with black or grey backgrounds, respectively. Positions of the conserved ATP-binding and catalytically active sites are highlighted with red and blue lines, respectively. B. The amino acid sequences of the *C. elegans* (PRK-1 and PRK-2), *Danio rerio* (PIM-1, PIM-2 and PIM-3), *Xenopus laevis* (PIM-1 and PIM-3), *Mus musculus* (PIM-1, PIM-2 and PIM-3) and *Homo sapiens* (PIM-1, PIM-2 and PIM-3) were used to draw a phylogenetic tree of PIM orthologs and paralogs in distinct species (www.phylogeny.fr). C. Radioactive kinase assays were performed to analyse the *in vitro* ability of mouse PIM-1 and *C. elegans* PRK-2 to phosphorylate themselves or NFATc1 in the absence (−) or presence (+) of 10 μM DHPCC-9. The intensities of phosphorylated proteins are shown in the upper panel and the total amounts of proteins in the lower panel. Shown are also the relative levels of phosphorylation of kinases and their substrates in control- (100%) versus drug-treated samples in this dataset representating three independent experiments.

A phylogenetic tree (Fig. 1B) shows the evolutionary distance between PIM orthologs and paralogs in several species, including also the zebrafish *Danio rerio* and the African clawed frog *Xenopus laevis*. In mammalian species and in frogs, there is more homology between each PIM ortholog than between the distinct family members, PIM-1, PIM-2 and PIM-3. By contrast, the *C. elegans* and *D. rerio* orthologs are all significantly separated from the other species. Interestingly and maybe also surprisingly, no obvious PIM orthologs have been found in the fruit fly *Drosophila melanogaster* (Peter Gallant, University of Würzburg, Germany, personal communication).

To demonstrate that the *C. elegans* PRKs are functional orthologs for mammalian PIM kinases, we took mouse PIM-1 and *Ce* PRK-2 proteins as representatives of each family and analysed for their activities by radioactive *in vitro* kinase assays. As expected and shown in Fig. 1C, both PIM-1 and PRK-2 were able to autophosphorylate themselves. More interestingly, also PRK-2 was able to phosphorylate the well-known PIM substrates NFATc1, BAD and RUNX3 (Fig. 1C and data not shown), suggesting that PIMs and PRKs are true orthologs sharing similar preferences for their target sites. Part of the samples were pretreated with the pyrrolocarbazole carbaldehyde DHPCC-9, which has been shown to efficiently and selectively inhibit catalytic activities of all three mammalian PIM kinases both *in vitro* (Akué-Gédu et al., 2009), in cell-based assays (Santio et al., 2010) and under *in vivo* conditions in mice (Santio et al., 2015) or chicken eggs (Santio et al., 2016a). At the 10 μM concentration used, DHPCC-9 was able to inhibit the activities of not only PIM-1, but also PRK-2 (Fig. 1C), indicating that the ATP-binding pockets of PIMs and PRKs are conserved enough to allow binding of the ATP-competitive drug. Even though the inhibition of PRK-2 was not as efficient as of PIM-1, it was sufficient to prompt us to use DHPCC-9 as a tool to investigate the physiological functions of PIM-related kinases in *C. elegans*.

### PIM inhibitors specifically suppress attractive olfactory responses via AWC^ON^ neurons

We have previously reported that mammalian PIM kinases are expressed in sensory organs such as the olfactory epithelium (Eichmann et al., 2000). Since PRK-1, and possibly also PRK-2, are expressed in *C. elegans* head neurons, we wanted to investigate whether they are involved in regulation of sensory functions, such as olfaction. Before carrying out behavioral assays, we experimentally determined that the maximal tolerated dosage of the PIM inhibitor DHPCC-9 in *C. elegans* is about 200 μM (data not shown). Then we synchronized wild-type animals, exposed populations of day one adults to 10 to 200 μM concentrations of DHPCC-9 for 120 min, and carried out chemotaxis assays to AWC-sensed attractive odorants for another 120 min. At tested dilutions of odorants, control animals vigorously responded to butanone, benzaldehyde and isoamyl alcohol (IAA) with chemotaxis indices (CI) of about 0.9, while 2,3-pentanedione was slighly less attractive with CI of 0.6 (Fig. 2A). More interestingly, animals pre-exposed to increasing concentrations of DHPCC-9 consistently displayed a significant dose-dependent decrease in their chemotaxis to butanone. The inhibitory effects of DHPCC-9 on the chemotactic behaviour became evident already at the lowest 10 μM concentration, but were most prominent at the maximal dose of 200 μM, where animals were moving randomly to all directions, as if they did not sense the presence of the odorant. There was a statistically significant reduction (about 20%) in chemotaxis to benzaldehyde in DHPCC-9-treated animals as compared to controls, while hardly any effects were observed by DHPCC-9 on chemotaxis to IAA or 2,3-pentanedione. It is known that butanone is sensed by AWC^ON^ neurons, 2,3-pentanedione by AWC^off^ neurons and benzaldehyde and IAA by both (Troemel et al., 1999; Wes and Bargmann, 2001). Thus, our data suggested that inhibition of PRK activity by DHPCC-9 specifically and asymmetrically suppresses olfactory sensing via AWC^ON^ neurons.

**Figure 2.**
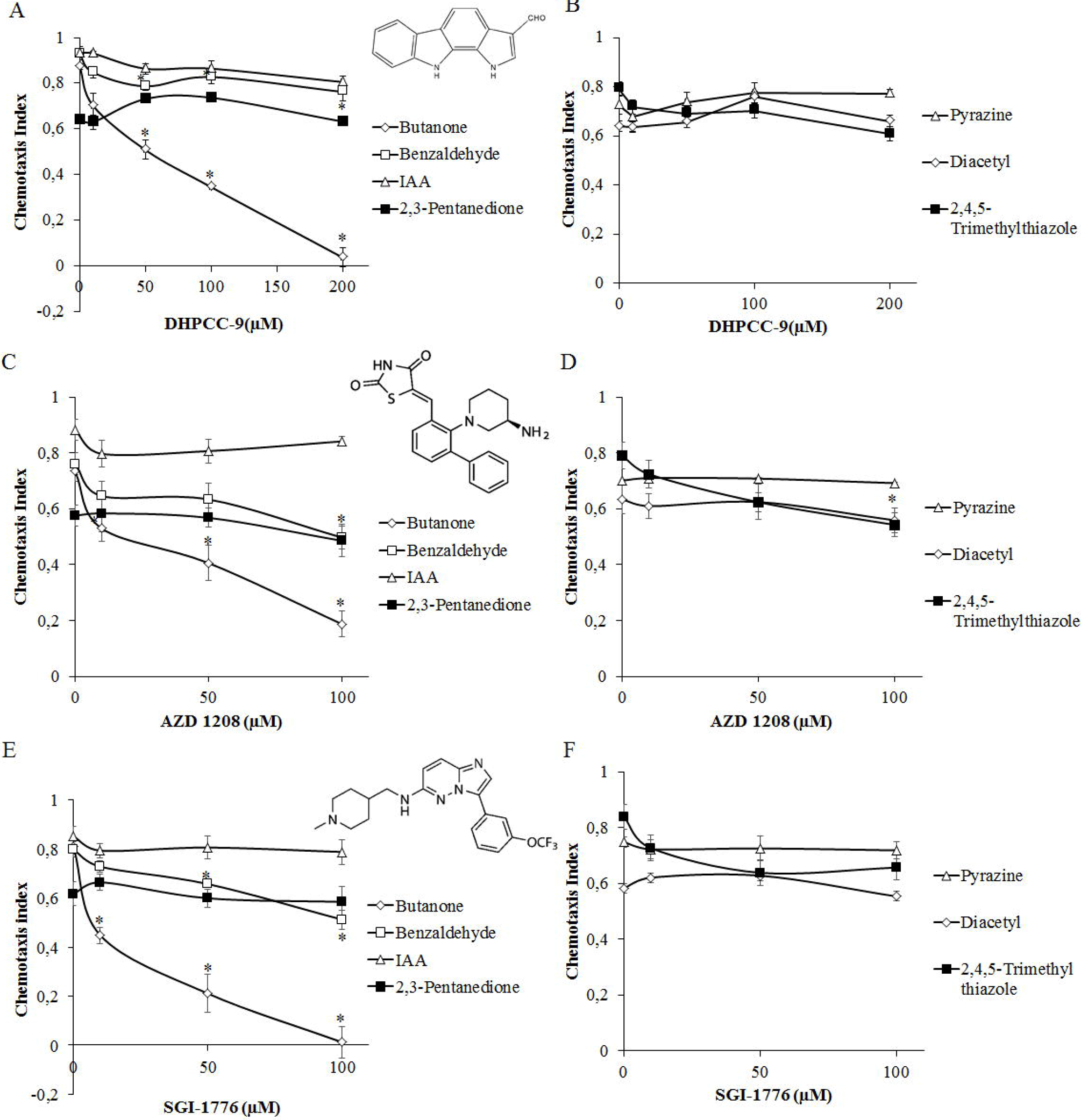
PIM inhibitors specifically suppress olfactory sensing via AWC^ON^ neurons. Synchronized day one adult animals were exposed to indicated concentrations of PIM inhibitors DHPCC-9 (A, B), AZD1208 (C, D) or SGI-1776 (E, F) for 120 min. Chemotaxis assays with odorants were performed for another 120 min. Chemical structures of the PIM inhibitors are shown in between the dose-dependent chemotactic indices of animals in response to AWC odorants (A, C and E) butanone (1:1000, 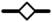), benzaldehyde (1:200,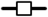 isoamyl alcohol (IAA, 1:100,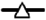), and 2,3-pentanedione (1:10000,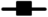) or AWA odorants (B, D and F) diacetyl (1:1000,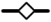), pyrazine (10 mg/ml, 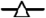) and 2,4,5-trimethylthiazole (1:1000, 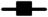). Each data point represents the means of at least six independent experiments with 150-200 animals per assay. Error bars indicate SEM and star (*) statistically significant differences (P value < 0.01) between control- and drug-exposed animals.

To examine whether DHPCC-9 affects the olfactory functions of also AWA neurons, we performed chemotaxis assays with AWA-sensed attractive odorants pyrazine, diacetyl and 2,4,5-trimethylthiazole. No significant suppression was observed for pyrazine or diacetyl by DHPCC-9, but the response to 2,4,5-trimethylthiazole, which can be sensed also by AWC neurons (Bargmann et al., 1993), was slightly reduced (about 15%) as compared to controls (Fig 2B).

To confirm that the observed effects of DHPCC-9 were dependent on its ability to inhibit endogenous PRK activity and were not just due to some unspecific off-target effects, we repeated the chemotaxis analyses by using two structurally unrelated PIM inhibitors, the imidazopyridazine SGI-1776 (Mumenthaler et al., 2009) and the thiazolidinedione AZD-1208 (Keeton et al., 2014). AZD-1208 efficiently targets all three PIM family members and is more selective than SGI-1776, which in addition to PIM-1 and PIM-3 also inhibits FLT-3 and haspin. These two inhibitors appeared to be more toxic than DHPCC-9, so animals were exposed to maximally 100 μM concentrations of them. When animals pre-exposed to either AZD-1208 or SGI-1776 were assayed for their chemotactic responses to attractive odorants sensed by either AWA or AWC neurons, very similar dose-dependent data were obtained as with DHPCC-9 (Fig. 2C-F). Again, chemotaxis to the AWC^ON^-sensed butanone was significantly suppressed by both AZD-1208 and SGI-1776. Chemotaxis to benzaldehyde and 2,4,5-trimethylthiazole was also reduced (about 20%) as compared to controls, while no significant effects were observed in animals treated with AWC^OFF^-sensed 2,3-pentanedione or AWA-sensed pyrazine and diacetyl. Overall, our data strongly suggested that the observed suppressive effects of all the drugs on olfactory sensations via AWC^ON^ neurons were due to their shared ability to inhibit activities of the PIM-related kinases.

### Inhibition of butanone sensing is reversible and dependent on odorant dosage

To determine whether or not the effects of the PIM inhibitors on butanone sensing were reversible, animals exposed for 120 min to 200 μM DHPCC-9 or 100 μM SGI-1776 were allowed to recover for up to 180 min before assaying their chemotaxis to butanone. As shown in Fig. 3A, drug-exposed animals quickly regained their ability to sense butanone, with complete recovery within 120 min or 180 min after exposure to DHPCC-9 or SGI-1776, respectively (Fig. 3B). These results indicate that the suppressive effects of the PIM inhibitors on olfactory sensations are reversible. Furthermore, the rapid inhibitory responses as well as recovery times suggest that the PIM inhibitors have direct, although as yet uncharacterized effects on olfactory signaling in the AWC^ON^ neurons.

**Figure 3.**
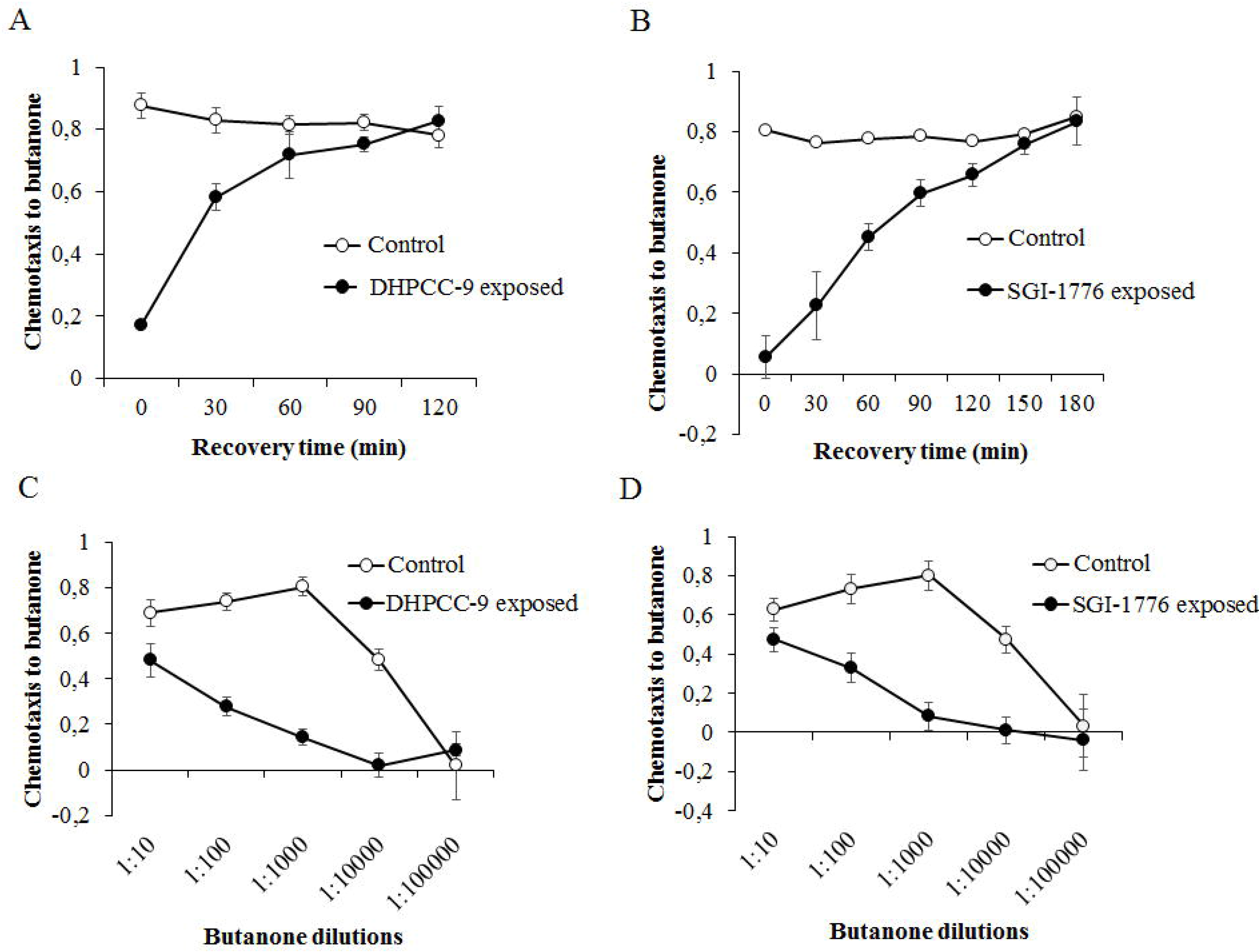
Inhibition of butanone sensing is reversible and dependent on odorant dosage. Animals exposed for 120 min to 200 μM DHPCC-9 (A and C) or 100 μM SGI-1776 (B and D) were assayed for chemotaxis to butanone (1:1000) after indicated intervals of recovery times (A and B) or to indicated dilutions of butanone (C and D). Open circles indicate chemotactic indices from control-exposed populations and closed circles from drug-exposed populations. Each data point represents the mean and SEM of four independent experiments with 150-200 animals per assay.

To investigate the effects of the PIM inhibitors on butanone sensing in more detail, control-and drug-treated animals were assayed for their chemotaxis to a wide range of butanone dilutions. As shown in Fig. 3C and D, high concentrations of butanone were able to overcome the inhibitory effects of both DHPCC-9 and SGI-1776, whereas at lower concentrations, the chemotactic behaviour of the animals was lost even without inhibition. Thus, these results confirmed that the 1:1000 dilution used in our other assays had been optimal to reveal the suppressive effects of the PIM inhibitors on butanone sensing.

### PIM inhibitors also repress repulsive olfactory responses via AWB neurons

To determine whether the PRKs are involved in the sensation of volatile repellants by AWB neurons, animals exposed for 120 min to increasing concentrations of DHPCC-9 (50, 100 and 200 μM) or SGI-1776 (50 and 100 μM) were tested for their ability to sense 1-octanol and to move away from it. After 120 min incubation, the avoidance indices were calculated. As shown in Fig. 4, more than half of the control-treated animals tried to avoid 1-octanol. By contrast, the avoidance responses were significantly and dose-dependently reduced in the drug-exposed populations, suggesting that the PIM inhibitors are able to diminish olfactory sensing also towards repellants via AWB neurons.

**Figure 4.**
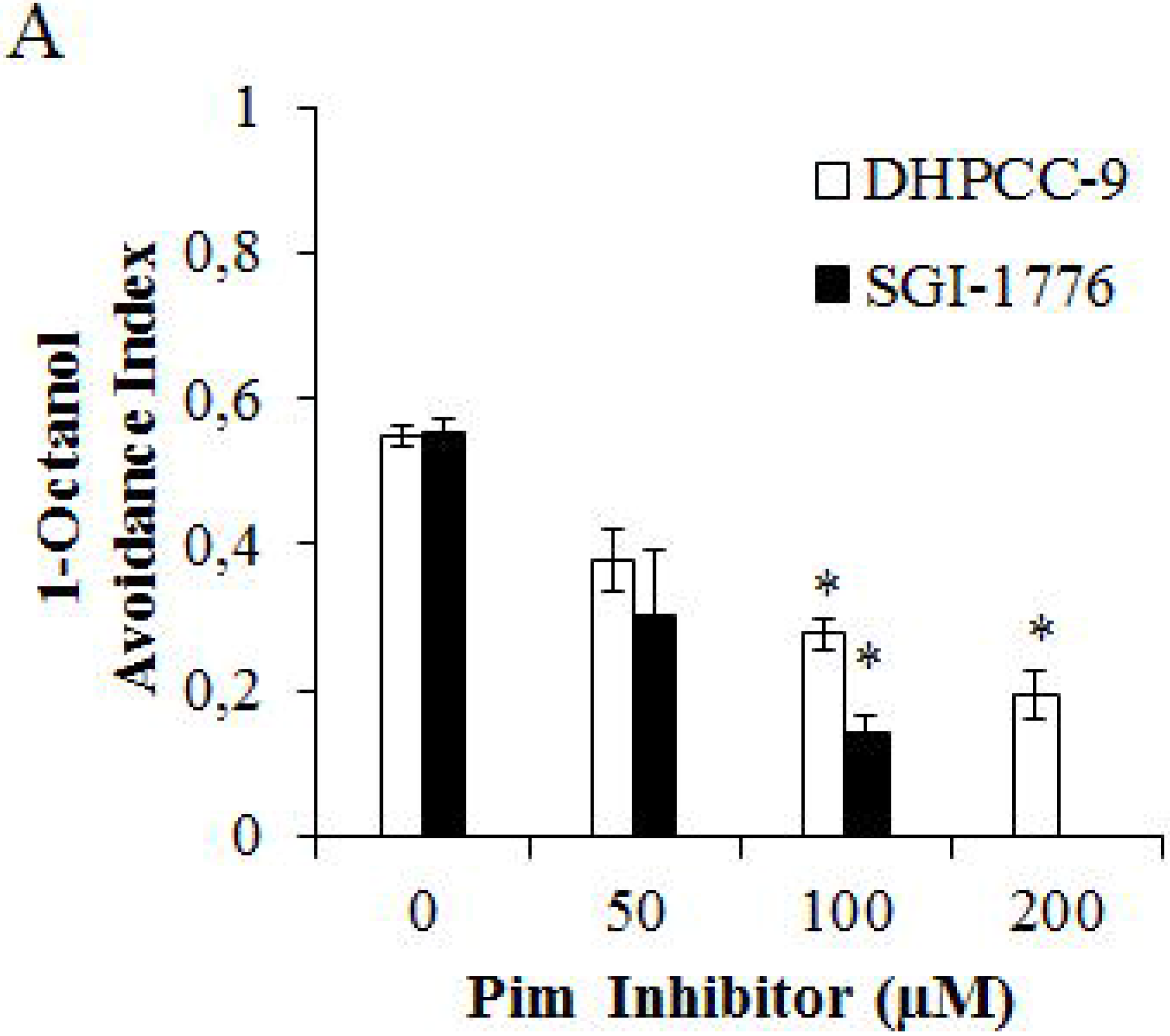
PIM inhibitors suppress sensing of olfactory repellants via AWB neurons. Animals were treated for 120 min with indicated concentrations of DHPCC-9 or SGI-1776. Aversion assays with 1-octanol (1:100) were performed for another 120 min. Shown are dose-dependent avoidance indices. Each data point represents the means of at least four independent experiments with 100-150 animals per assay. Error bars indicate SEM and star (*) statistically significant differences (P value < 0.01) between control- and drug-exposed animals.

### PIM inhibitors do not affect gustatory sensations

We next examined whether PRKs are essential also for the attractive and aversive gustatory responses by the ASE and ASH neurons, respectively. Similarly to AWC neurons, also the ASE neurons are functionally asymmetric: the ASER (right) neuron preferentially detects potassium ions, while the ASEL (left) neuron detects sodium ions (Pierce-Shimomura et al., 2001). When we placed animals exposed to 200 μM DHPCC-9 or 100 μM SGI-1776 on plates containing drops of either 2.5 M NaCl or KCl, and calculated the chemotactic indices after 120 min incubation, more than half of the animals were attracted to either salt, but the PIM inhibitors did not significantly interfere with these responses as compared to controls (Fig. 5A).

**Figure 5.**
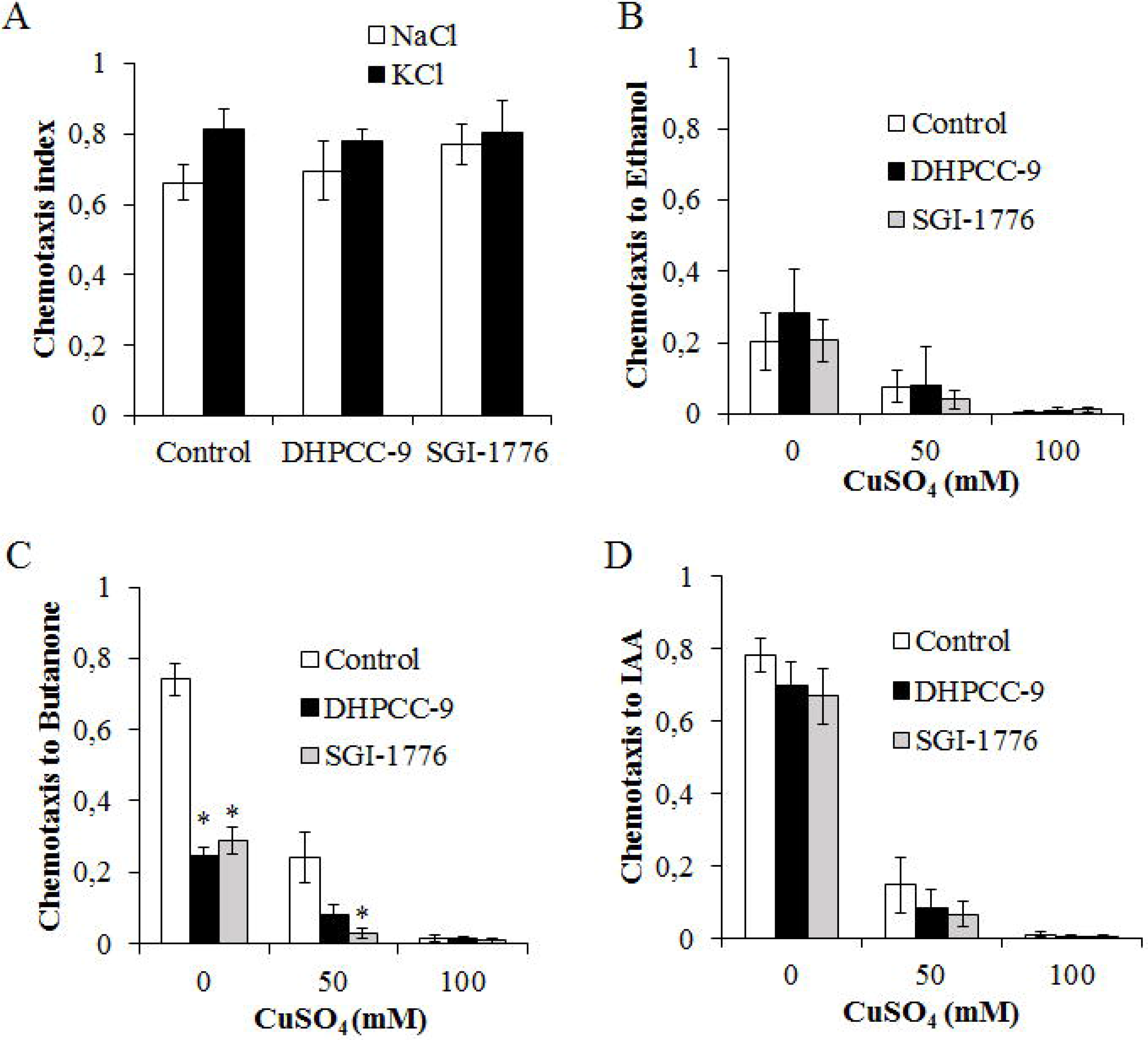
PIM inhibitors do not affect responses to gustatory attractants or repellants. Animals were treated for 120 min with 200 μM DHPCC-9 or 100 μM SGI-1776 and assayed for responses to gustatory attractants (A) or repellents (B, C and D). (A) Shown are chemotactic indices of control-or drug-exposed animals to 2.5 M NaCl or 2.5M KCl. Each data point represents the mean and SEM of three independent experiments with 150-200 animals per assay. (B) Indicated concentrations of CuSO_4_ were pipetted across the midlines of the assay plates. Control-or drug-exposed animals were placed on one side and drops of ethanol (B), butanone (C) or IAA (D) to the other side. After 120 min of incubation, chemotactic indices to the odorants were calculated. Each data point represents the mean and SEM of four independent experiments with 200-250 animals per assay. (*) statistically significant differences (P value < 0.01) between control- and drug-exposed animals.

To investigate the repulsive responses by ASH neurons, midlines of distilled water or CuSO_4_ (50 or 100 mM) were drawn onto midlines of assay plates. Control or drug-exposed animals were placed on one side of the midline, whereas aliquots of ethanol, butanone or IAA were dropped on the other side. After 120 min incubation, chemotactic indices were calculated. As shown in Fig. 5B, the initially mild chemotaxis towards the neutral odorant ethanol was further reduced by increasing concentrations of CuSO_4_ in the midline, while the PIM inhibitors did not have significant effects. Both butanone (Fig. 5C) and IAA (Fig. 5D) efficiently attracted the animals to cross the midline of water, but the chemotactic responses were strongly diminished by the presence of 50 mM CuSO_4_ and completely blocked by 100 mM CuSO_4_. As observed also in our olfactory assays (Fig. 2A, C and E), exposures to either 200 μM DHPCC-9 or 100 μM SGI-1776 efficiently inhibited the positive chemotaxis towards butanone, but had no major effects on IAA sensing.

### PRK-1 is expressed in amphid and phasmid neurons

To confirm that PRK kinases are present in the amphid neurons, whose activities are suppressed by the PIM inhibitors, we wanted to analyse the PRK expression patterns in more detail. Unfortunately there are no antibodies available against PRKs to allow direct detection of these proteins. However, using the *prk-1::GFP* reporter strain, we were able to confirm previous data on *prk-1* promoter-driven GFP expression in the intestine, as well as head and tails neurons (Fig 6A). A more detailed investigation revealed that the reporter is expressed in a few amphid neurons, possibly including AWB and AWC (Fig. 6B and 6C) and in one phasmid neuron (Fig. 6D). These results support our observations from the olfactory assays and suggest that PRK-1, but possibly also PRK-2 is present in AWB and AWC neurons to regulate the olfactory responses to volatile attractants and repellants.

**Figure 6.**
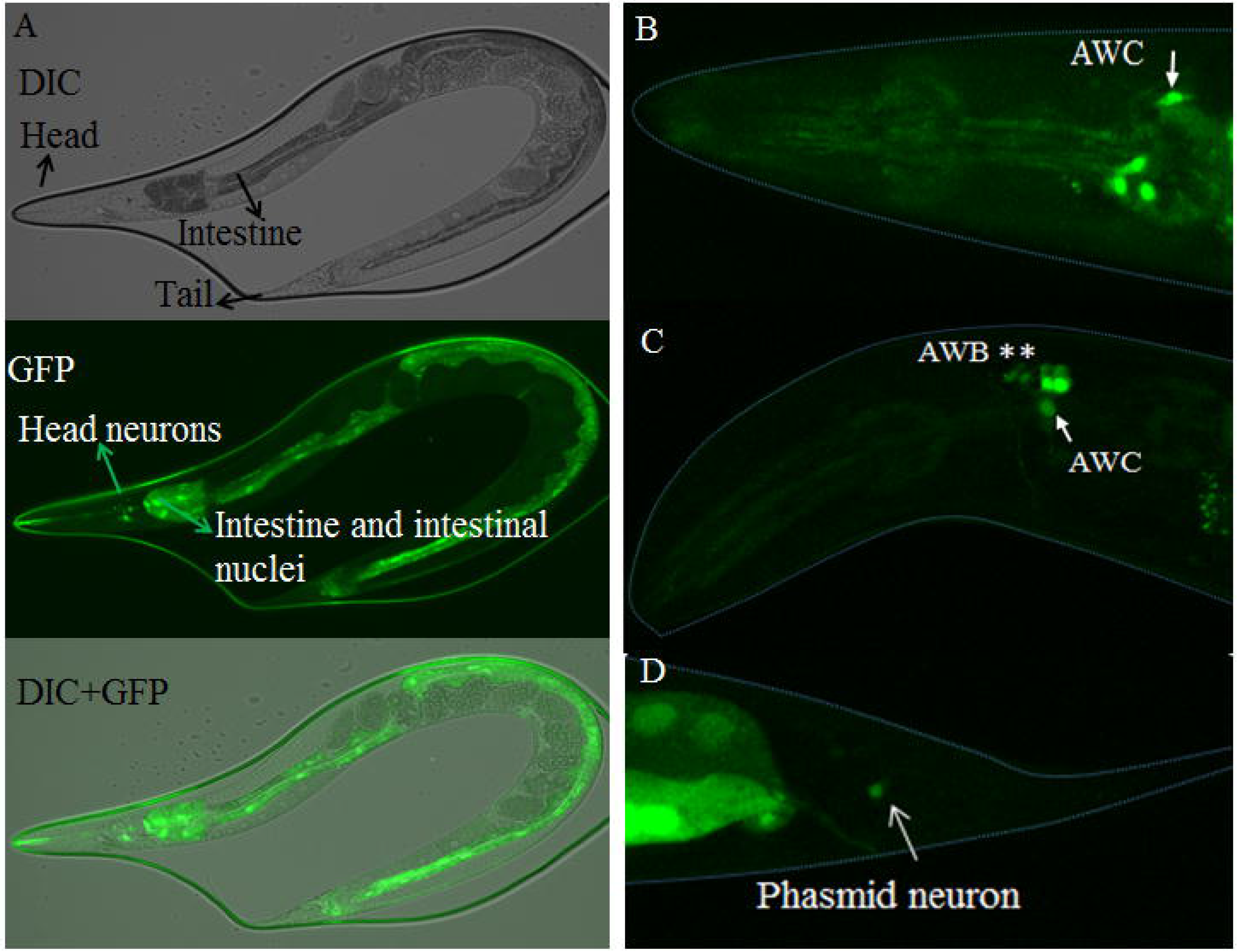
Expression pattern of PRK-1. (A) Lateral view of an adult from the *prk-1*∷GFP strain expressing GFP in amphid and phasmid neurons, and in the intestine. (B-D) Confocal images showing *prk-1*-driven GFP expression in AWB(*), AWC(→) and one phasmid neuron.

## Discussion

In this study, we have partially characterized the functional roles of *C. elegans* PRK family members in regulation of chemosensation and shown that they are true orthologs for the three mammalian PIM family kinases. Based on our *in vitro* kinase data, PRK-2 is autophosphorylated and able to target several known PIM substrates. Since the amino acid sequences within the kinase domains of PIM and PRKs are highly conserved, especially at the ATP-binding site and the catalytically active site, it is highly likely that also PRK-1 shares substrates with PRK-2 and PIM family members. However, the sequences of different lengths outside the kinase domains do not show any homology between the mammalian and *C. elegans* proteins, and are not conserved between PRK-1 and PRK-2, either. Thus, these sequences may allow more individual interactions of distinct PRK family members with other proteins.

The catalytic activity of the mammalian PIM kinases can be blocked by selective ATP-competitive inhibitors, such as DHPCC-9. Crystallization studies have revealed that within the ATP-binding pocket of murine PIM-1, DHPCC-9 forms hydrophobic contacts to the highly polar region including Lys67 (Akué-Gédu et al., 2009), which in *C. elegans* PRK-1 corresponds to Lys87 and in PRK-2 to Lys60. Our data from *in vitro* kinase assays indicate that the structures of the ATP-binding pockets of PIM orthologs are so well conserved from mammals to *C. elegans* that it is possible to use PIM-selective inhibitors as tools to study physiological functions of PRKS. However, in the absence of structural data on the interactions of PIM inhibitors with PRKS, we cannot completely rule out any off-target effects, which is why we found it important to use three structurally unrelated compounds in our assays, including SGI-1776 and AZD-1208.

According to our data from chemotaxis assays, all the three tested PIM-selective inhibitors have very similar effects on *C. elegans* chemosensation. All of them can in a dose-dependent fashion block chemotaxis towards butanone sensed by the AWC^ON^ neurons, with complete suppression seen already at sublethal concentrations of the inhibitors. In addition, small but significant suppressive effects are observed towards benzaldehyde sensed by both AWC^ON^ and AWC^OFF^ neurons. By contrast, responses to 2,3-pentanedione sensed by AWC^OFF^ neurons or pyrazine and diacetyl sensed by AWA neurons remain unaffected, whereas some suppression is detected in chemotaxis to trimethylthiazole sensed by both AWA and AWC neurons. These results strongly suggest that PRK activity is essential for the olfactory functions of AWC^ON^ neurons, but is dispensable for sensations via AWA or AWC^OFF^ neurons. Furthermore, the suppressive effects of PIM inhibitors on butanone sensing are reversible and dependent also on the dosage of the attractant, as high butanone concentrations can overcome the suppression, whereas lower concentrations reduce the chemotaxis in both control-and drug-exposed animals.

As demonstrated in the presence of the repellant octanol, PIM inhibitors can block also aversive responses to volatile compounds via AWB neurons. By contrast, responses to water-soluble gustatory attractants such as NaCl and KCl via ASE neurons are not affected. Combinatory assays with both olfactory and gustatory cues revealed that chemotaxis towards attractive odorants is efficiently blocked by a barrier of water-soluble metal compounds such as copper sensed via ASH neurons. While PIM inhibitors do not interfere with such aversive responses, they selectively reduce AWC^ON^ -dependent sensing of butanone also in this setting.

In conclusion, our data indicate that PRKs are involved in regulation of olfactory attraction via AWC^ON^ neurons and repulsion via AWB neurons, but do not affect gustatory sensations via ASE and ASH neurons. These results are further supported by our observations on the *prk-1* promoter-driven reporter expression in amphid neurons, possibly including both AWB and AWC. Here it should be noted that AWC^ON^ and AWC^OFF^ neurons are structurally similar, but differ from each other by randomly selected expression of the G protein-coupled seven transmembrane receptor (STR-2) on either right or left side of the animals, resulting in functional differences in the ability of these neurons to detect olfactory cues (Wes and Bargmann, 2001). Since PIM inhibitors disrupt sensory functions of only AWC^ON^ neurons expressing STR-2, the putative connection between PRKs and STR-2 should be analysed in more detail. In addition, it should be determined whether both PRK family members are equally essential for the regulation of olfactory sensations.

The present data are well in line with our previous observation that during mouse embryogenesis, PIM kinases are expressed in the olfactory epithelium, but not in any gustatory organs (Eichmann et al., 2000). Thus, it would be interesting to test whether mice lacking *pim* genes or treated with PIM inhibitors have problems in olfactory sensing. Knockout mice without any of the three *pim* family genes are viable, but smaller than mice with at least one *pim* allele (Mikkers et al., 2004), which may be explained by our previous data on the essential role of PIM kinases in supporting cytokine responses (Rainio et al., 2002; Aho et al., 2004). Furthermore, the triple knockout mice are fertile, but hard to breed (Anton Berns, National Cancer Institute, Amsterdam, the Netherlands, personal communication), which according to our data might be explained by problems in pheromone sensing. In case PIM inhibitors were able to reduce also human olfactory responses, this phenomenon could even be used as a biomarker for the efficacy of PIM-targeted therapies that are currently being developed against several types of hematological or solid cancers.

In developing mouse embryos, PIM kinases are expressed also in the neural retina (Eichmann et al., 2000), suggesting that PIM inhibition may negatively affect not only olfaction, but also vision. Indeed, evidence for that has already been obtained from studies with *D. rerio* zebrafish embryos (Yin et al., 2012). Using an antibody against human PIM-1, low and high PIM expression was observed in the neural retina at 3 and 5 days post fertilization, respectively. Furthermore, genetic or pharmacological inhibition of PIM expression or activity significantly diminished the visual functions of the embryos without affecting retinal morphology. Thus, these data together with our observations suggest that PIM-related kinases selectively affect sensory functions in an evolutionarily conserved fashion. However, further studies are still needed to identify the relevant PIM substrates involved and to dissect the molecular mechanisms in more detail.

## ACKNOWLEDGEMENTS

These studies were financed by the Academy of Finland (grants 297700 and 287040 to P.J.K, 297776 to C.I.H). *C. elegans* strains were provided by the Caenorhabditis Genetics Center, which is funded by NIH Office of Research Infrastructure Programs (P40 0D010440). The Koskinen and Holmberg group members, P. Gallant and A. Berns are acknowledged for helpful discussions, and M. Nonet for reagents.

## REFERENCES

Aho, T.L., Sandholm, J., Peltola, K.J., Mankonen, H.P., Lilly, M., and Koskinen, P.J. (2004). Pim-1 kinase promotes inactivation of the pro-apoptotic Bad protein by phosphorylating it on the Ser112 gatekeeper site. FEBS Lett. 571, 43–49.

Aho, T.L., Sandholm, J., Peltola, K.J., Ito, Y., Koskinen, P.J. (2006). Pim-1 kinase phosphorylates RUNX family transcription factors and enhances their activity. BMC Cell Biol. 7, 21.

Akué-Gédu, R., Rossignol, E., Azzaro, S., Knapp, S., Filippakopoulos, P., Bullock, A.N., Bain, J., Cohen, P., Prudhomme, M., Anizon, F., and Moreau, P. (2009). Synthesis, kinase inhibitory potencies, and in vitro antiproliferative evaluation of new Pim kinase inhibitors. J. Med. Chem. 52, 6369–6381.

Bargmann, C.I. (2006). Chemosensation in C. elegans. WormBook: the online review of C elegans biology, 1–29.

Bargmann, C.I., Hartwieg, E., and Horvitz, H.R. (1993). Odorant-selective genes and neurons mediate olfaction in C. elegans. Cell 74, 515–527.

Brault, L., Gasser, C., Bracher, F., Huber, K., Knapp, S., and Schwaller, J. (2010). PIM serine/threonine kinases in the pathogenesis and therapy of hematologic malignancies and solid cancers. Haematologica 95, 1004–1015.

Brenner, S. (1974). The genetics of Caenorhabditis elegans. Genetics 77, 71–94.

Eichmann, A., Yuan, L., Breant, C., Alitalo, K., and Koskinen, P.J. (2000). Developmental expression of Pim kinases suggests functions also outside of the hematopoietic system. Oncogene 19, 1215–1224.

Keeton, E.K., McEachern, K., Dillman, K.S., Palakurthi, S., Cao, Y., Grondine, M.R., Kaur, S., Wang, S., Chen, Y., Wu, A., Shen, M., Gibbons, F.D., Lamb, M.L., Zheng, X., Stone, R.D., DeAngelo, D.J., Platania, L.C., Dakin, L.A., Chen, H., Lyne, P.D., and Huszar, D. (2014). AZD1208, a potent and selective pan-Pim kinase inhibitor, demonstrates efficacy in preclinical models of acute myeloid leukemia. Blood 123, 905–913.

Kiriazis, A., Vahakoski, R.L., Santio, N.M., Arnaudova, R., Eerola, S.K., Rainio, E.M., Aumuller, I.B., Yli-Kauhaluoma, J., and Koskinen, P.J. (2013). Tricyclic Benzo[cd]azulenes selectively inhibit activities of Pim kinases and restrict growth of Epstein-Barr virus-transformed cells. PLoS One 8, e55409.

McKay S.J., Johnsen R., Khattra J., Asano J., Baillie D.L., Chan S., Dube N., Fang L., Goszczynski B., Ha E., Halfnight E., Hollebakken R., Huang P., Hung K., Jensen V., Jones S.J., Kai H., Li D., Mah A., Marra M., McGhee J., Newbury R., Pouzyrev A., Riddle D.L., Sonnhammer E., Tian H., Tu D., Tyson J.R., Vatcher G., Warner A., Wong K., Zhao Z., Moerman D.G. (2003). Gene expression profiling of cells, tissues, and developmental stages of the nematode C. elegans. Cold Spring Harb. Symp. Quant. Biol. 68, 159–169.

Mikkers, H., Nawijn, M., Allen, J., Brouwers, C., Verhoeven, E., Jonkers, J., and Berns, A. (2004). Mice deficient for all PIM kinases display reduced body size and impaired responses to hematopoietic growth factors. Mol. Cell. Biol. 24, 6104–6115.

Mumenthaler, S.M., Ng, P.Y., Hodge, A., Bearss, D., Berk, G., Kanekal, S., Redkar, S., Taverna, P., Agus, D.B., and Jain, A. (2009). Pharmacologic inhibition of Pim kinases alters prostate cancer cell growth and resensitizes chemoresistant cells to taxanes. Mol. Cancer Ther. 8, 2882–2893.

Nawijn, M.C., Alendar, A., and Berns, A. (2011). For better or for worse: the role of Pim oncogenes in tumorigenesis. Nat. Rev. 11, 23–34.

Pierce-Shimomura, J.T., Faumont, S., Gaston, M.R., Pearson, B.J., and Lockery, S.R. (2001). The homeobox gene lim-6 is required for distinct chemosensory representations in C. elegans. Nature 410, 694–698.

Rainio, E.M., Sandholm, J., and Koskinen, P.J. (2002). Cutting edge: Transcriptional activity of NFATc1 is enhanced by the Pim-1 kinase. J Immunol 168, 1524–1527.

Santio, N.M., Eerola, S.K., Paatero, I., Yli-Kauhaluoma, J., Anizon, F., Moreau, P., Tuomela, J., Harkonen, P., and Koskinen, P.J. (2015). Pim Kinases promote migration and metastatic growth of prostate cancer xenografts. PLoS One 10 (6), e0130340.

Santio, N.M., and Koskinen, P.J. (2017). PIM kinases: From survival factors to regulators of cell motility. Int. J. Biochem. Cell Biol. 93, 74–85.

Santio, N.M., Landor, S.K., Vahtera, L., Yla-Pelto, J., Paloniemi, E., Imanishi, S.Y., Corthals, G., Varjosalo, M., Manoharan, G.B., Uri, A., Lendahl, U., Sahlgren, C., and Koskinen, P.J. (2016a). Phosphorylation of Notch1 by Pim kinases promotes oncogenic signaling in breast and prostate cancer cells. Oncotarget 7, 43220–43238.

Santio, N.M., Salmela, M., Arola, H., Eerola, S.K., Heino, J., Rainio, E.M., and Koskinen, P.J. (2016b). The PIM1 kinase promotes prostate cancer cell migration and adhesion via multiple signalling pathways. Exp. Cell Res. 342, 113–124.

Santio, N.M., Vahakoski, R.L., Rainio, E.M., Sandholm, J.A., Virtanen, S.S., Prudhomme, M., Anizon, F., Moreau, P., and Koskinen, P.J. (2010). Pim-selective inhibitor DHPCC-9 reveals Pim kinases as potent stimulators of cancer cell migration and invasion. Mol. Cancer 9, 279.

Troemel, E.R., Kimmel, B.E., and Bargmann, C.I. (1997). Reprogramming chemotaxis responses: sensory neurons define olfactory preferences in C. elegans. Cell 91, 161–169.

Troemel, E.R., Sagasti, A., and Bargmann, C.I. (1999). Lateral signaling mediated by axon contact and calcium entry regulates asymmetric odorant receptor expression in C. elegans. Cell 99, 387–398.

Wes, P.D., and Bargmann, C.I. (2001). C. elegans odour discrimination requires asymmetric diversity in olfactory neurons. Nature 410, 698–701.

Wicks, S.R., de Vries, C.J., van Luenen, H.G., and Plasterk, R.H. (2000). CHE-3, a cytosolic dynein heavy chain, is required for sensory cilia structure and function in Caenorhabditis elegans. Dev. Biol. 221, 295–307.

Yan, B., Zemskova, M., Holder, S., Chin, V., Kraft, A., Koskinen, P.J., and Lilly, M. (2003). The PIM-2 kinase phosphorylates BAD on serine 112 and reverses BAD-induced cell death. J. Biol. Chem. 278, 45358–45367.

Yin, J., Shine, L., Raycroft, F., Deeti, S., Reynolds, A., Ackerman, K.M., Glaviano, A., O’Farrell, S., O’Leary, O., Kilty, C., Kennedy, C., McLoughlin, S., Rice, M., Russell, E., Higgins, D.G., Hyde, D.R., and Kennedy, B.N. (2012). Inhibition of the Pim1 oncogene results in diminished visual function. PLoS One 7, e52177.

Yu, S., Avery, L., Baude, E., and Garbers, D.L. (1997). Guanylyl cyclase expression in specific sensory neurons: a new family of chemosensory receptors. Proc. Natl. Acad. Sci. U.S.A. 94, 3384–3387.

Zheng, Q., Schaefer, A.M., and Nonet, M.L. (2011). Regulation of C. elegans presynaptic differentiation and neurite branching via a novel signaling pathway initiated by SAM-10. Development 138, 87–96.

